# Mathematical connection between short telomere induced senescence calculation and mortality rate data

**DOI:** 10.1101/2020.06.15.152348

**Authors:** Jerry B. Torrance, Steve Goldband

## Abstract

The last 20 years have seen a surge in scientific activity and promising results in the study of aging and longevity. Many researchers have focused on telomeres, which are composed of a series of TTAGGG repeat nucleotide sequences at the ends of each chromosome. Measurements of the length of these telomere strands show that they decrease in length with increasing age, leading many authors to propose that when the length of these telomere strands decreases sufficiently, the cells enter into a state of replicative senescence, eventually leading to disease and death. These ideas are supported by evidence that short telomere length is correlated with increased mortality. In this paper, we extend this idea to make an actual calculation of the predicted mortality rate caused by short telomere length induced senescence (STLIS). We derive a simple equation for the mathematical relationship between telomere length and mortality rate. Using only 3 parameters based on telomere length measurement data of Canadians, we have calculated both the magnitude and the age dependence of the mortality rate, for both men and women. We show that these calculated data are in good quantitative agreement with the actual number of Canadians that die. This agreement provides strong evidence (but not proof) that the mechanism of STLIS plays an important role in the major diseases of aging (e.g., cardiovascular disease, many cancers, and diabetes mellitus) which dominate human mortality. This result represents significant progress in our understanding the factors behind the cause of aging.

## Measurements of telomere lengths

Many mechanisms have been proposed as the cause of aging [1,2]. In this paper, we will focus on a model based on telomeres, whose lengths has been shown [3–6] correlate with the mortality of a number of diseases of aging. Since the initial work on telomeres in the 1970’s, more than a hundred studies [6–9] have measured the telomere length as a function of age for a large number of individuals, using different techniques. As an example, Fig 1 shows the data of Aubert et al [7] for male Canadians, using the Flow FISH technique.

**Fig 1:**
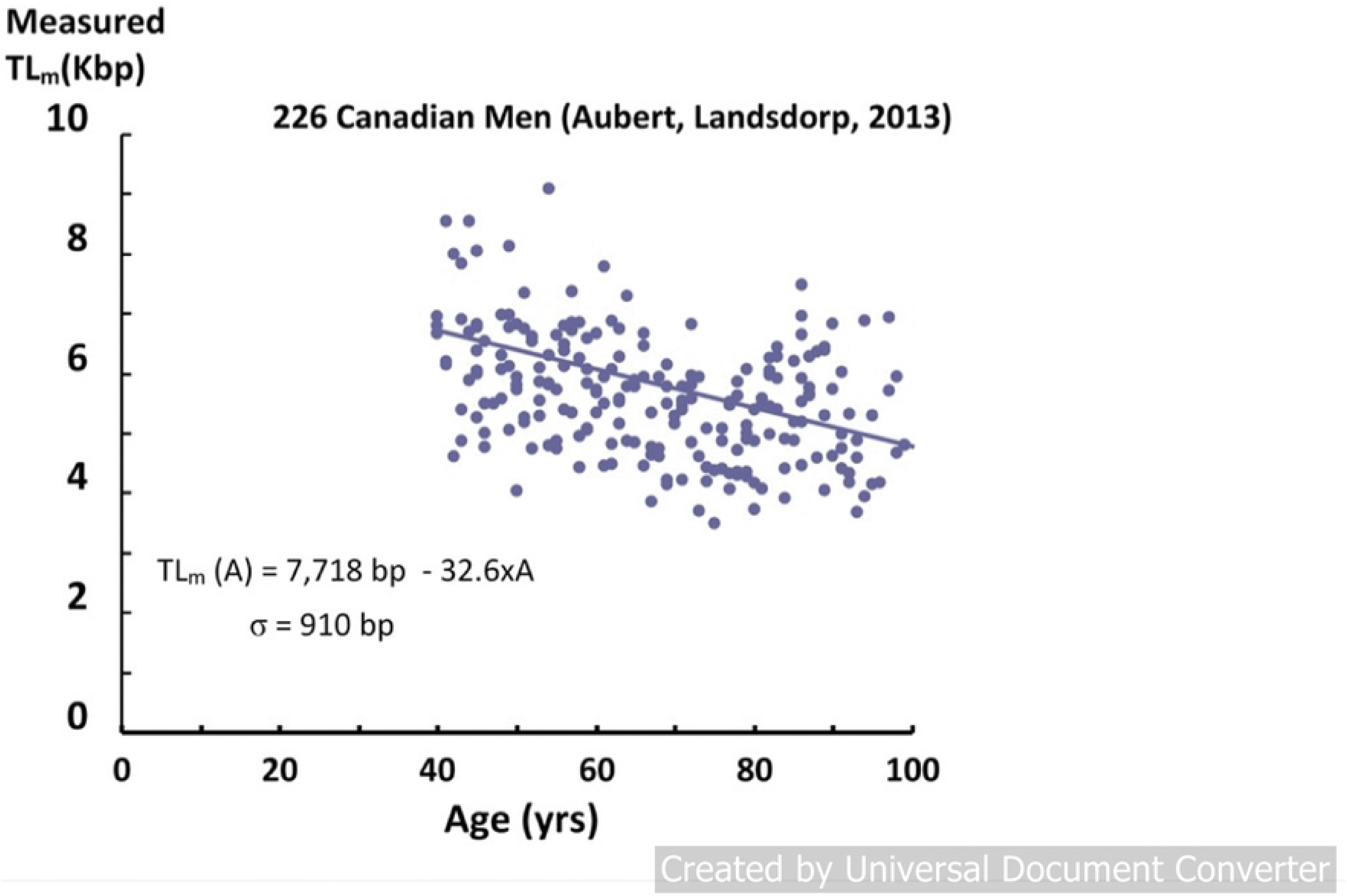
The measured telomere length (in Kbp, thousands of base pairs) using the Flow FISH technique is plotted as a function of age for 226 Canadian males, along with a linear fit to the data.

Each data point corresponds to the measured “average” telomere length for one individual. The first feature of the data is that different individuals with the same age have telomere lengths that differ by as much as a factor of 2. These differences have been attributed to differences in heredity, lifestyles, exposures to inflammation and oxidation, telomerase activity, as well as stress. This distribution of lengths among these individuals is presumed to be a normal distribution, similar to the distribution in heights among different individuals, for example. With this assumption, we can calculate the standard deviation (**σ**) by fitting the telomere length data (using STEYX in Excel, Table 1). We find that **σ** is very large (890-900 bp) and varies little with age (A>40), as observed in other measurements [8]. This standard deviation measures the width of the distribution of measured telomere lengths for different individuals having the same age.

**Table 1:**
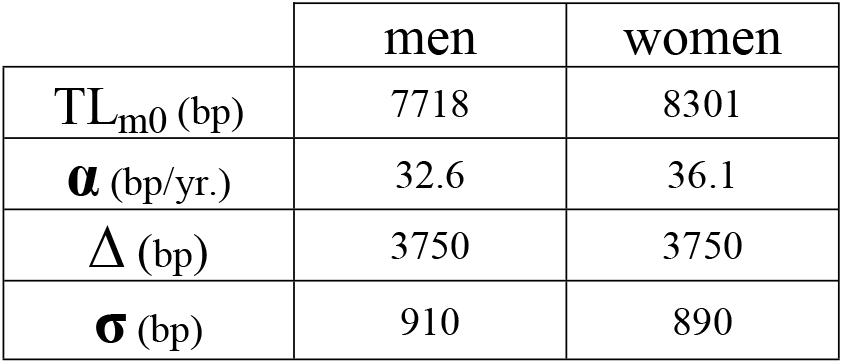
Values of 4 telomere measurement-based parameters.

|  | men | women |
| --- | --- | --- |
| $TL_{m0}$ (bp) | 7718 | 8301 |
| $\alpha$ (bp/yr.) | 32.6 | 36.1 |
| $\Delta$ (bp) | 3750 | 3750 |
| $\sigma$ (bp) | 910 | 890 |

The second feature of the data is that with increasing age, there is a clear decrease in measured telomere lengths. This decrease is due, in part, to cellular replication, in which the length of the telomeres shortens with each division. Changes with age in the factors mentioned above also contribute (both positively and negatively) to a drop in telomere length (TL). This decrease is observed in more than a hundred experiments [6–10], where the measured telomere length (TL_m_) is well approximated by a linear decrease with age (A):

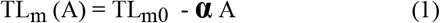

In which the parameters TL_m0_ and **α** are constants. For the Canadian data in Fig 1, these parameters (obtained by fitting the telomere data) are shown in Table 1 for both men and women.

### Short telomere length induced senescence

Since the discovery of telomeres, hundreds of papers have been published reporting measurements of telomere lengths and their correlation with lifestyle factors and with longevity, building the evidence for using telomere length as a cellular marker of aging [3–6]. Furthermore, many authors [11–15] have proposed that aging may be caused by shortened telomere lengths inducing senescence (STLIS). The general idea behind this model is that when the telomere length decreases with age and becomes “sufficiently short”, the cells stop replicating (Hayflick limit) and go into a state of senescence. In this state, they begin secreting inflammatory chemicals (SASP) into the cell, which induce disease and subsequent death. But the measured telomere lengths (Fig 2) are far from going to zero, even for individuals of 100 years old.

**Fig 2:**
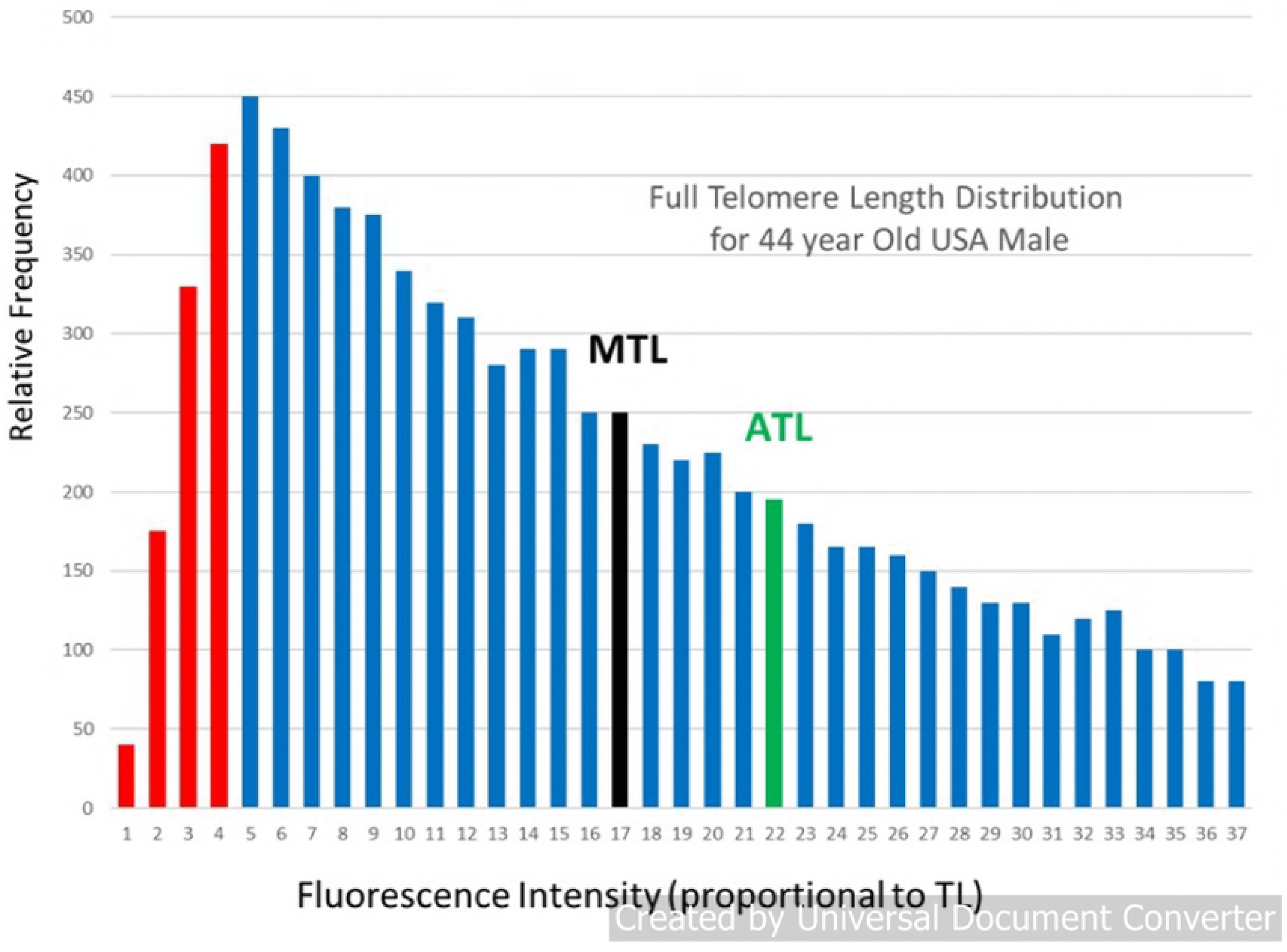
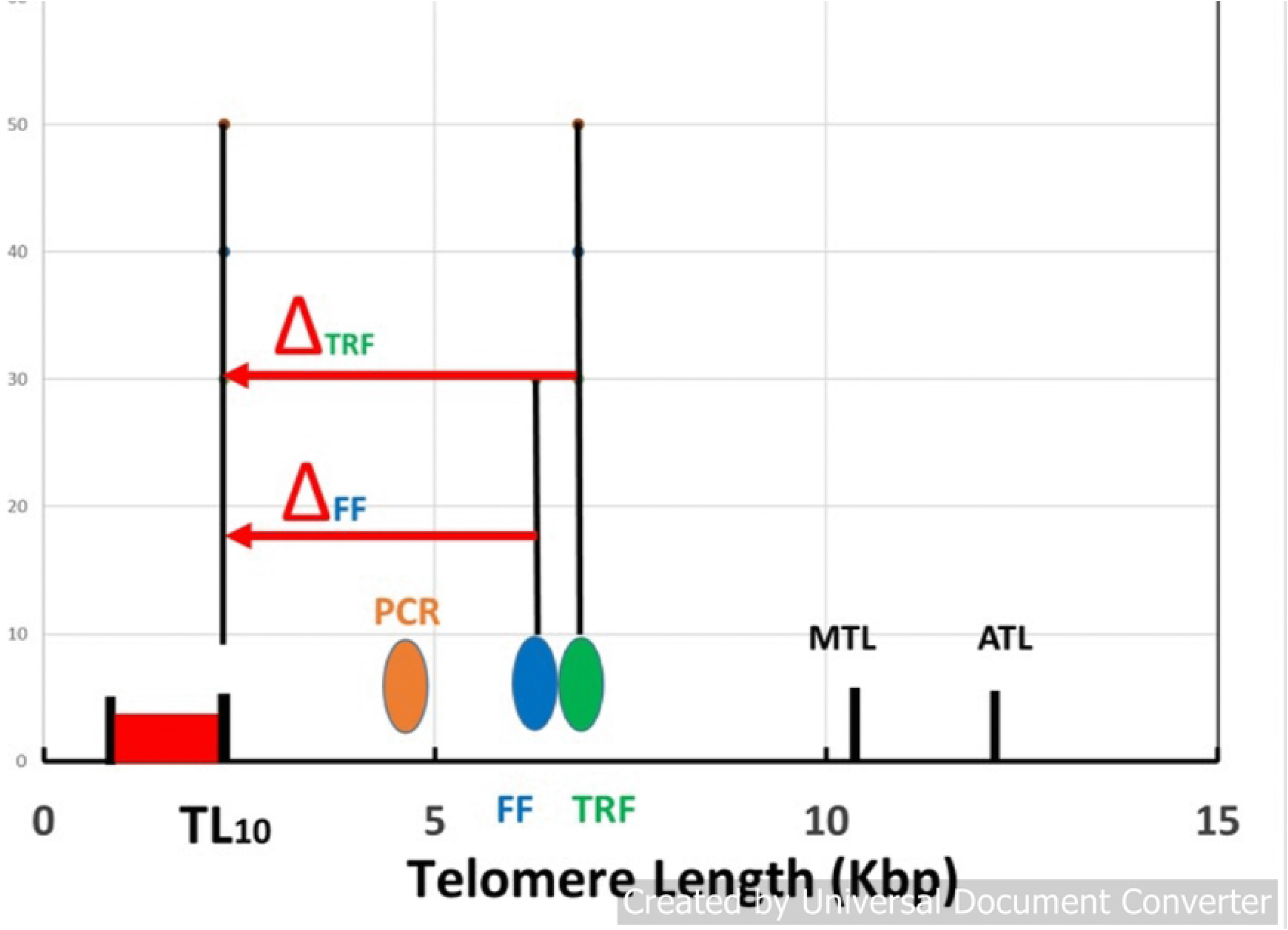
(a) Relative frequency of telomeres as a function of fluorescence intensity (proportional to telomere length) for a 44 year-old USA male, (b) A schematic figure, comparing the telomere lengths measured by different techniques. The bubbles represent the single “average” telomere length measured by the PCR, FF, and TRF techniques, compared with the full telomere length distribution, including zero length, the 10^th^ Percentile (TL_10_), the Median (MTL) and Average (ATL) lengths and even more at higher lengths, all included in the HT-Q-FISH measurement.

This apparent conflict is related to the fact that each individual in Fig 2 is represented as a single point, with a single measurement of his/her telomere length. However, in reality, each individual has a large number of telomeres and they have a wide range of telomere lengths. This can be shown by using a more sophisticated measurement technique, in which the full distribution of telomere lengths for an individual can be measured. The High Throughput Q-FISH (Quantitative Fluorescence In-Situ Hybridization) technique is used in laboratories of only a very few universities [16] and one company [17]. An example of this measurement is shown in Fig 2a for a 44 year-old USA male with average measured telomere length [18].

This typical individual has a very asymmetric distribution of widely different telomere lengths (Fig 2a), ranging from several hundred base pairs to tens of thousands [17]. Because of the asymmetry, the Median and Average Telomere Lengths (MTL and ATL) are much longer than the peak of the distribution. Also shown are the critical lowest length telomeres, those below the 10th percentile, which are marked in red.

In Fig 2(b), we compare the results from different measurement techniques [19]. The Median and Average Telomere length (MTL and ATL) from Fig 2a are shown from HT Q-FISH measurements, along with the shortest 10% of the telomere lengths (in red). Other techniques measure only a single number for this wide distribution of telomere lengths, which is some kind of weighted average of the full distribution. Examples from measurements using the “Flow FISH” (FF) technique for the average 44-year-old male Canadian [7] and using the TRF (Terminal Restriction Fragment) technique for the average 44-year-old Danish male [8] are represented by the data points shown as the blue and green bubbles in Fig 2b at 6,284 bp and 6,768 bp, respectively. Results using the popular PCR (Polymerase Chain Reaction) technique vary considerably. The orange bubble shown in Fig 2b is the data point for an average 44-year-old Danish male, using this technique [6]. The last three techniques measure only a single “average” length, which lies near the peak of the distribution for the individual (Fig 2a) and which is substantially lower than the actual median (MTL) (10,500 bp) and the average (ATL) (12,400 bp) values (a consequence of the very skewed distribution in Fig 2a). More significantly, they are significantly higher than the lowest (red) TL which are those involved in senescence. These different “telomere lengths” measured by different techniques have caused some confusion in the literature. In the rest of this paper, we shall use the term “measured” telomere length, TL_m_, to refer to the values of the “average” telomere lengths measured by these latter techniques.

The models of short telomere length induced senescence (STLIS) predict that when the lengths of the critically lowest telomeres (in red) become “sufficiently short”, the cells in this individual will senesce and diseases of aging will begin. However, these models do not specify what is actually meant by “sufficiently short”. We shall arbitrarily estimate that cellular senescence will have been induced when the lowest 10% of the telomere lengths have decreased to zero. A critical parameter for this model is then the difference in telomere length between the **measured** length of the individual and the length of his 10 percentile telomeres, TL_10_, in Fig 2b. This difference we call Δ which is defined as Δ = TL_m_ - TL_10_. The STLIS model then can be described with the use of Fig 2: as this individual ages, his entire telomere length distribution (Fig 2a) will shift to lower lengths, making the important assumption (for simplicity) that it will maintain approximately the same shape [17]. Thus, both TL_m_ and TL_10_ also shift together to lower lengths, with the difference in TL between them (Δ) remaining constant. The point when the TL_10_ has decreased to zero (and 10% of the telomeres are below zero) is the point at which the cells will have started to senesce. (Note that this will occur long before the **measured** telomere length (TL_m_) (the bubbles in Fig 2b) approach zero.)

The parameter Δ will be different for each technique. For Flow-FISH measurements, for example, the value of Δ is estimated from Fig 2b to be near 4,000 bp, whereas for TRF it would be closer to 4,500 bp. For HT-Q-FISH it would be nearer to 8,000 bp (using MTL as the measured value). A large number of authors have discussed the importance of the critically short length telomeres [16]. Using TRF, the Aviv group [20] has introduced the related concept of a “telomere brink”: when the measured telomere length decreases and approaches the “telomere brink”, there is a high risk of subsequent death. The brink was estimated to be about 5,000 bp, similar to the value of 4,500 bp, estimated above.

### Calculation of mortality rate due to STLIS

In order to calculate the mortality at any given age, we need to look at populations of individuals having that age. In Fig 3, we show the number of individuals aged 40, 60 and 80 years old calculated from using a normal distribution and the telomere data in Table 1, obtained from fitting the data in Fig 1. These are plotted vs. their TL_10_ since it is these, shortest telomeres that are going to senesce and induce eventual mortality, according to the STLIS model.

**Fig 3:**
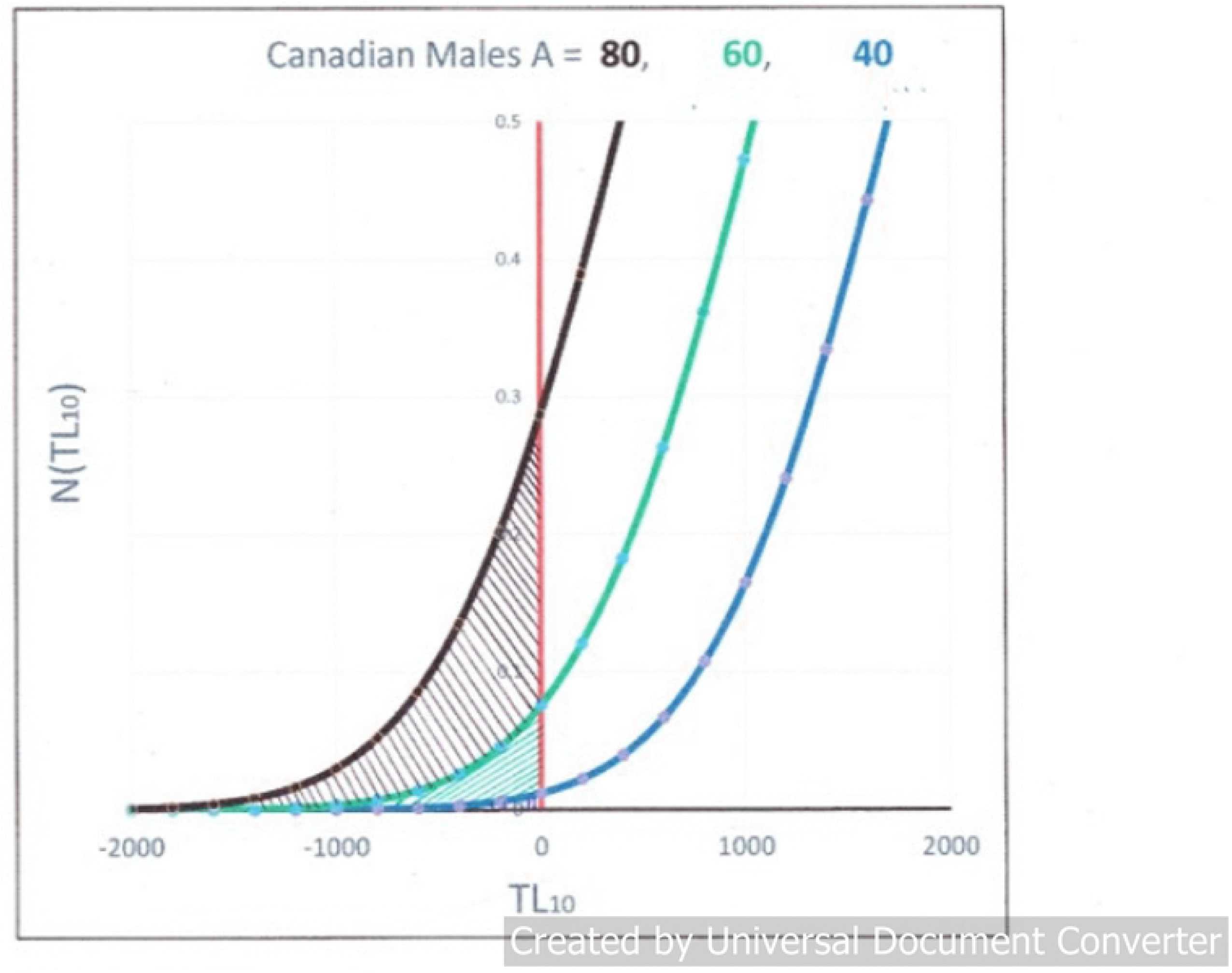
Number of individuals in the population/100K vs. their measured telomere length for ages=40, 60, and 80 years. The number of people expected to have died is the integrated number of people whose TL_10_ is negative, i.e., the hashed area under the curves for different ages.

At age A, the people dying are those individuals who have the shortest measured telomere length relative to the rest of the population and consequently their shortest individual telomere lengths have gone to zero. The total number of these people predicted to have died at a certain age A is the mortality rate, M_R_(A), and is equal to the total number of individuals for whom 10% of their telomere lengths have gone to zero, i.e., those whose TL_10_ is less than 0. This is obtained by integrating the area under the curves in Fig 3 for negative TL_10_, i.e., the hashed areas under the left side of the distribution. In this figure, one can see how dramatically this number increases with increasing age.

Mathematically, we can write this integration as:

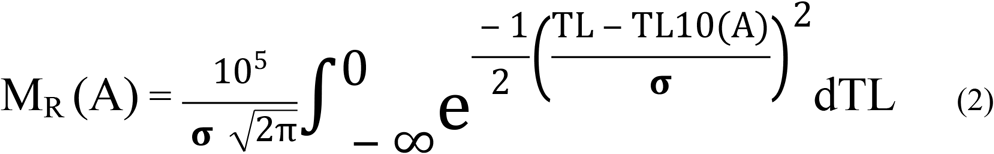

(which may be calculated in Excel by M_R_(A) = 10^5^ x NORMDIST (0, (TL_m0_ - Δ - **α** A), **σ**, TRUE).

Notice that the mortality rate is dependent basically on only 3 telomere parameters: the standard deviation (**σ**), the slope from Fig 1 (**α**), and the **difference** (TL_m0_ - Δ). For a specific age, A, we obtain M_R_(A) using the NORMDIST function to calculate the number of them under the left side of the normal distribution curves of Fig 3 for negative TL_10_, i.e., the hashed areas.

Compared with earlier correlations [3–6] between shorter telomere lengths and higher mortality, the present calculation (Equation 2) gives a complete mathematical relation between telomere length and mortality, for the STLIS model. Using Equation 2 and three telomere parameters (2 of which are measured and 1 estimated), we calculate the mortality rate, M_R_(A), as a function of age for the STLIS model.

### Actual mortality rate data and agreement with calculated values

Since the telomere data in Fig 1 and Table 1 are for Canadians, we show the mortality rate data [21] for Canadian women (solid red lines) and men (solid blue lines) in Fig 4.

**Fig 4:**
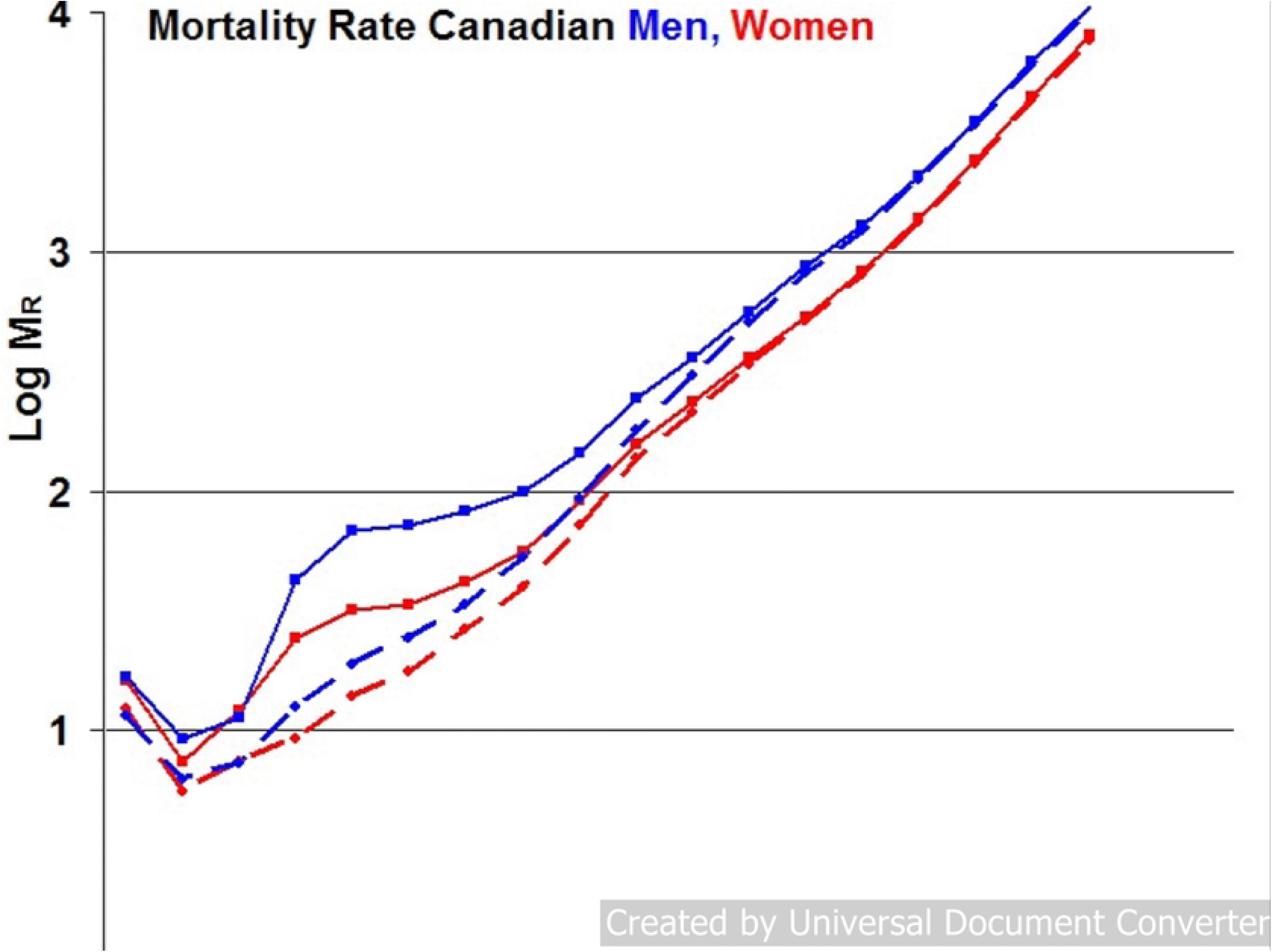
The solid lines are the logarithm of the age-dependent total mortality rates per 10^5^ population for Canadian men (blue) and women (red) in 2013. The dashed lines are the same data, after subtracting the mortality due to Accidents, Suicides, and Assaults.

The characteristic feature of these and other mortality data is the strong, exponential increase in the mortality rate (M_R_) with increasing age, especially above 40 years, as recognized by Gompertz [22]. This increase is attributed to “Diseases of Aging”, in which age is the dominant risk factor, such as cardiovascular disease, cancers, and Alzheimer’s. At younger ages, however, other types of mortality are significant as well. For example, the Canadian mortality database includes deaths due to accidents, suicides, and assaults. We have subtracted the known mortality of these causes from the total mortality data and obtained the two dashed lines in Fig 4. Even though the bump in the original data near age 25 is removed, there remains some mortality below age 40 due to infant mortality, infections, childhood cancers, and other diseases not related to aging. In order to minimize the effect of these, we shall concentrate on ages greater than 40 years, where the mortality data are dominated by diseases of aging.

In Fig 5, we compare the actual mortality data (dashed lines from Fig 4) for A>40 (after subtracting the Accidents-Suicides-Assaults contribution) to the mortality rate data predicted by Equation 2 and the telomere data in Table 1 (solid lines), where data for men are in blue and women in red.

**Fig 5:**
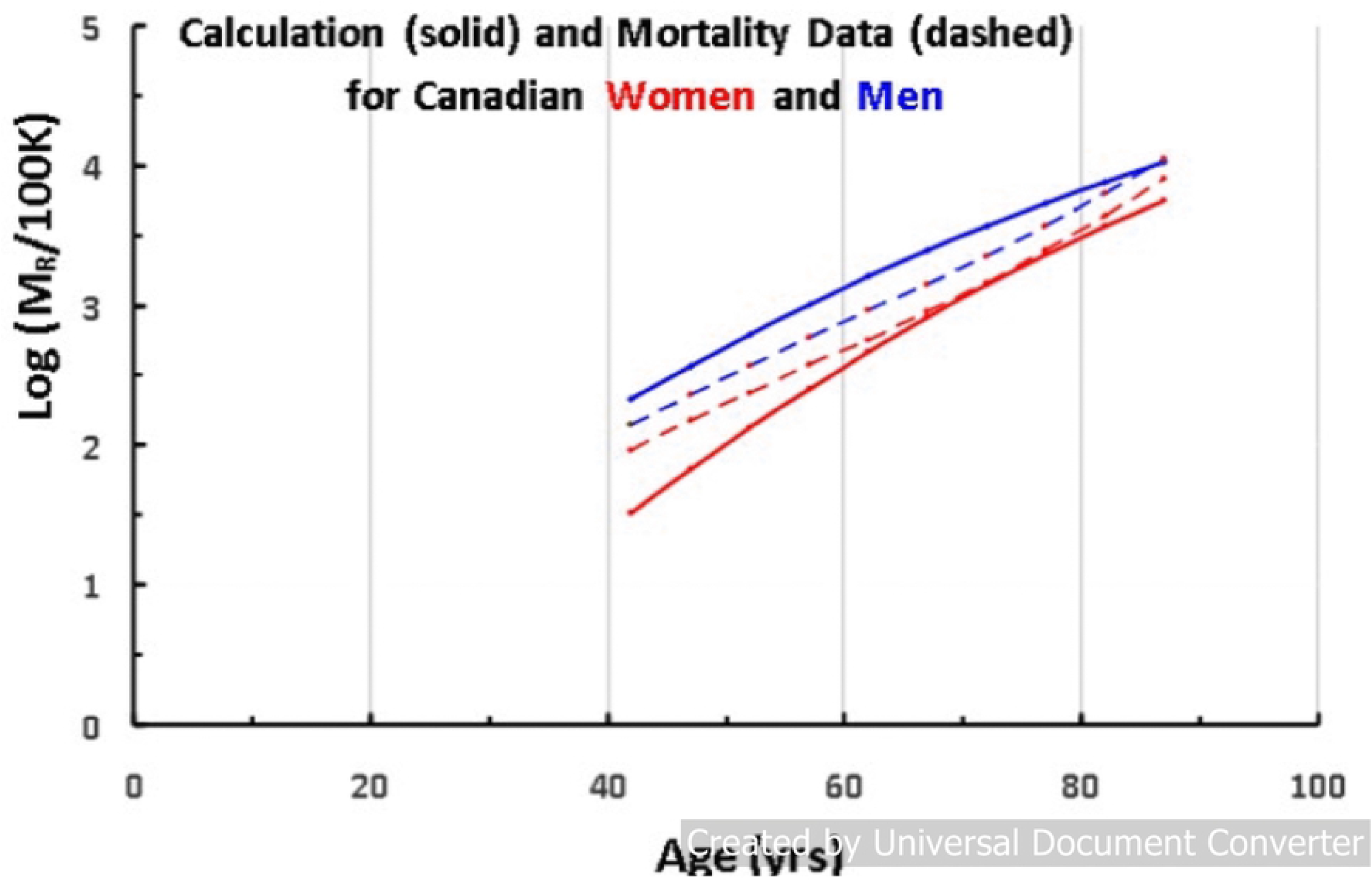
The log of the mortality rate (per 100K) vs. age for men (red) and women (blue), comparing the calculated data (solid lines) with actual data (dashed lines) for Canadians (2013).

For a better agreement, we used Δ = 3,750 bp as the value for both men and women, instead of our initial estimate of 4,000bp (Fig 2). (Note that using a different value for Δ for men and for women, we would obtain an even better fit to the data, but it would add another parameter.)

The results of the calculation are in good agreement with both the magnitude and the age dependence of the mortality, for both men and for women. Such an agreement should perhaps not have been expected, considering the fact that the data have extra mortality due to non-aging diseases and the fact that the model describes the onset of the disease, while the data refers to death (often delayed by treatment). In addition, this simple model is calculating the mortality for such very different diseases, as different as cancer and cardiovascular disease and Alzheimer’s.

In conclusion, it has been proposed [11–15] that short telomere lengths induce senescence (STLIS) and that this mechanism is an important cause of the diseases of aging. Using this model and 3 parameters based on only telomere measurements, we have calculated the magnitude of the all-cause mortality rate and its age dependence in good agreement with the actual number of people who die each year and their age dependence, for both men and women. This result provides strong evidence (but not proof) that the STLIS model plays an important role in most of the diseases of aging.

## Author Contributions

J.B.T. initiated the project and the calculations. S.G. contributed important ideas.

## Acknowledgements

We thank Life Length for sharing the telomere length distribution in Fig 2a. We acknowledge on-going important and provocative discussions with F. Shepardson. STEYX and NORMDIST are programs in Excel.

## References

1. Lopez-Otin C, Blasco MA, Partridge L, Serrano M, and Kroemer G. The hallmarks of aging. Cell. 2013; 153: 1194–1217.

2. “The Longevity Issue”, MIT Tech. Rev., 2019; Sept/Oct.

3. Cawthon RM, Smith KR, O’Brien E, Sivatchenko A, and Kerber RA, Association between telomere length in blood and mortality in people aged 60 years or older. Lancet. 2003; 361:393–395.

4. Epel ES, Merkin SS, Cawthon R, Blackburn EH, Adler NI, Pletcher MJ, et al. The rate of leukocyte telomere shortening predicts mortality from cardiovascular disease in elderly men, Aging (Albany NY). 2009; 1: 81–86.

5. Wang Q, Zhan Y, Pederson NL, Fang F, & Hagg S, Telomere Length and All-Cause Mortality: A Meta-Analysis, Ageing Res. Rev. 2018; 48: 11–21.

6. Rode L, Nordestgaard BG and Bojesen SE, Peripheral blood leukocyte telomere length and mortality among 64637 individuals from the general population, J. Natl. Cancer Inst. 2015; 107:djv074.

7. Aubert G, Baerlocher GM, Vulto I, Poon SS and Lansdorp PM, Collapse of telomere homeostasis in hematopoietic cells caused by heterozygous mutations in telomerase gene, PLoS Genetics 2012; 8:e1002696.

8. Factor-Litvak P, Susser E, Kezios K, McKeague I, Kark JD, Hoffman M, et al. Leukocyte telomere length in newborns: implications for the role of telomeres in human disease, Pediatrics 2017; 137:(4)e20153927.

9. Dalgard C, Benetos A, Verhulst S, Labat C, Kark JD, Christensen K, et al. Leukocyte telomere length dynamics in women and men: menopause vs age effects, Int.J.Epidemiol. 2015; 44: 1688–1695.

10. Muezzinler A, Zineddin AK, and Brenner H, A systematic review of leukocyte telomere length and age in adults, Ageing Res. Rev. 2013; 12: 509–519.

11. Bodnar, A.G. Ouellette M, Frolkis M, Holt SE, Chiu CP, Morin GB, et al. Extension of life-span by introduction of telomerase into normal human cells, Science 1988; 279: 349–352.

12. Harley CB, Futcher AB, and Greider CW, Telomeres shorten during ageing of human fibroblasts. Nature. 1990; 345: 458–460.

13. Andrews B, and Cornell J, “Telomere Lengthening: Curing all Diseases including Cancer and Aging”, Sierra Sciences, LLC, www.sierrasci.com

14. Fossel M, Blackburn G, and Woynarowski D, “The Immortality Edge: Realize the Secrets of Your Telomeres for a Longer, Healthier Life”, Wiley (2010).

15. Anderson R, Lagnado A, Maggiorani D, Walaszczyk A, Dookun E, Chapman J, et al. Length-independent telomere damage drives post-mitotic cardiomyocyte senescence, EMBO J. 2019; 38:e100492.

16. Vera E, and Blasco MA, Beyond average: potential for measurement of short telomeres, Aging 2012; 4: 379392.

17. Life Length; de Pedro N, Diez M, Garcia I, Garcia J, Otero L, Fernandez L, et al. P, Bio. Proced. Online, 2020; 22: 2–8.

18. Data provided by Life Length, Madrid.

19. Montpetit AJ, Alhareeri AA, Montpetit M, Starkweather AR, Elmore LW, Filler K, et al. Telomere length: a review of methods of measurement, Nurs. Res. 2014; 63: 289–299.

20. Steenstrup T, Kark JD, Verhulst S, Thinggaard M, Hjelmborg JVB, Dalgard C, et al., Telomeres and the natural lifespan limit in humans, Aging 2017; 9: 1130–1142.

21. Statistics Canada, www.stat.can.gc.ca

22. Gompertz B, On the nature of the function expressive of the law of human mortality, and on a new mode of determining the value of life contingencies, Phil. Trans. Royal Soc. London, Bio. Sci. 1825; 182: 513–585.

